# Evolution and Diversification Dynamics of Butterflies

**DOI:** 10.1101/2022.05.17.491528

**Authors:** Akito Y. Kawahara, Caroline Storer, Ana Paula S. Carvalho, David M. Plotkin, Fabien Condamine, Mariana P. Braga, Emily A. Ellis, Ryan A. St Laurent, Xuankun Li, Vijay Barve, Liming Cai, Chandra Earl, Paul B. Frandsen, Hannah L. Owens, Wendy A. Valencia-Montoya, Kwaku Aduse-Poku, Emmanuel F. A. Toussaint, Kelly M. Dexter, Tenzing Doleck, Amanda Markee, Rebeccah Messcher, Y-Lan Nguyen, Jade Aster T. Badon, Hugo A. Benítez, Michael F. Braby, Perry A. C. Buenavente, Wei-Ping Chan, Steve C. Collins, Richard A. Rabideau Childers, Even Dankowicz, Rod Eastwood, Zdenek F. Fric, Riley J. Gott, Jason P. W. Hall, Winnie Hallwachs, Nate B. Hardy, Rachel L. Hawkins Sipe, Alan Heath, Jomar D. Hinolan, Nicholas T. Homziak, Yu-Feng Hsu, Yutaka Inayoshi, Micael G.A. Itliong, Daniel H. Janzen, Ian J. Kitching, Krushnamegh Kunte, Gerardo Lamas, Michael J. Landis, Elise A. Larsen, Torben B. Larsen, Jing V. Leong, Vladimir Lukhtanov, Crystal A. Maier, Jose I. Martinez, Dino J. Martins, Kiyoshi Maruyama, Sarah C. Maunsell, Nicolás Oliveira Mega, Alexander Monastyrskii, Ana B. B. Morais, Chris J. Müller, Mark Arcebal K. Naive, Gregory Nielsen, Pablo Sebastián Padrón, Djunijanti Peggie, Helena Piccoli Romanowski, Szabolcs Sáfián, Motoki Saito, Stefan Schröder, Vaughn Shirey, Doug Soltis, Pamela Soltis, Andrei Sourakov, Gerard Talavera, Roger Vila, Petr Vlasanek, Houshuai Wang, Andrew D. Warren, Keith R. Willmott, Masaya Yago, Walter Jetz, Marta A. Jarzyna, Jesse W. Breinholt, Marianne Espeland, Leslie Ries, Robert P. Guralnick, Naomi E. Pierce, David J. Lohman

## Abstract

Butterflies are a diverse and charismatic insect group that are thought to have diversified via coevolution with plants and in response to dispersals following key geological events. These hypotheses have been poorly tested at the macroevolutionary scale because a comprehensive phylogenetic framework and datasets on global distributions and larval hosts of butterflies are lacking. We sequenced 391 genes from nearly 2,000 butterfly species to construct a new, phylogenomic tree of butterflies representing 92% of all genera and aggregated global distribution records and larval host datasets. We found that butterflies likely originated in what is now the Americas, ∼100 Ma, shortly before the Cretaceous Thermal Maximum, then crossed Beringia and diversified in the Paleotropics. The ancestor of modern butterflies likely fed on Fabaceae, and most extant families were present before the K/Pg extinction. The majority of butterfly dispersals occurred from the tropics (especially the Neotropics) to temperate zones, largely supporting a “cradle” pattern of diversification. Surprisingly, host breadth changes and shifts to novel host plants had only modest impacts.

## Background

Butterflies have captivated naturalists, scientists, and the general public for centuries, and they have played a central role in studies of speciation, community ecology, plant-insect interactions, mimicry, genetics, and conservation. Despite being the most studied group of insects, butterfly evolutionary history and drivers of their diversification are still poorly understood^1,2^. Due to over a century of efforts by amateur naturalists, butterflies are one of the few insect groups for which substantial trait data (e.g., geographic and host plant information) exist to test diversification hypotheses. Until now, these data have largely been scattered across the literature, museum collections, and local databases. Butterflies are thought to have diversified in relation to multiple abiotic and biotic factors, including adaptations to novel climates and species interactions, with geographic history and caterpillar-host interactions playing a major role^3^. However, these have not been studied because a synthesis of associated traits and a robust phylogenetic framework at the taxonomic scale needed to examine their evolution has not been available.

We sequenced 391 genes from nearly 2,000 butterfly species to construct a new, robust, phylogenomic tree of butterflies representing 92% of all genera, aggregated global distribution records, and assembled a comprehensive host association dataset. Using this tree, we infer the evolutionary timing, biogeographic history, and diversification patterns of butterflies. We address three long-standing questions related to butterfly evolution: 1) did butterflies originate in the northern (Laurasia) or southern (Gondwana) hemisphere?^4^, 2) what plants did the ancestor of butterflies feed on?^5^, and 3) were plants a major driver promoting butterfly diversification?^6^

## Results and Discussion

To elucidate patterns of global butterfly diversification in space and time, we used targeted exon capture ^7^ to assemble a dataset of 391 gene regions (161,166 nucleotides and 53,722 amino acids) from 2,244 butterfly species. The majority (1,914 specimens) of butterflies sampled were newly sequenced for this study, and represented all families, tribes, and 92% of recognized genera, with most samples coming from museum collections across the world (see Supplementary Materials). Phylogenomic trees inferred with nucleotides and amino acids were highly congruent, with strong support for the monophyly of all families and nearly all subfamilies with traditional branch support metrics (Table S1), multispecies coalescence (MSC) analyses (Table S1), and 4-cluster likelihood mapping (FcLM) (Table S2). Our results strongly support the need for revision of the classification of at least 32 butterfly tribes (25% of total) as currently circumscribed (Table S1).

We conducted 24 dating analyses using different fossil and secondary calibration schemes along with sensitivity analyses to assess the impact of analytical and sampling bias. Across analyses, our results revealed largely congruent timing of butterfly divergence events (Table S3), indicating that butterflies originated from nocturnal, herbivorous moth ancestors during the mid-Cretaceous (101.4 Ma, 102.5–100.0 Ma). This provides independent evidence for the Cretaceous origin recovered in previous butterfly phylogenies^2,8^. We conducted diversification rate analyses that used time-variable models and clade-heterogeneous models that are less prone to sampling bias (Supplementary Methods). Both approaches recovered similar patterns of diversification across butterflies, with increased rates in clades such as the diverse skipper tribe Eudamini and lycaenid subfamily Poritiinae (Fig. 1, Fig. S1).

**Fig. 1.**
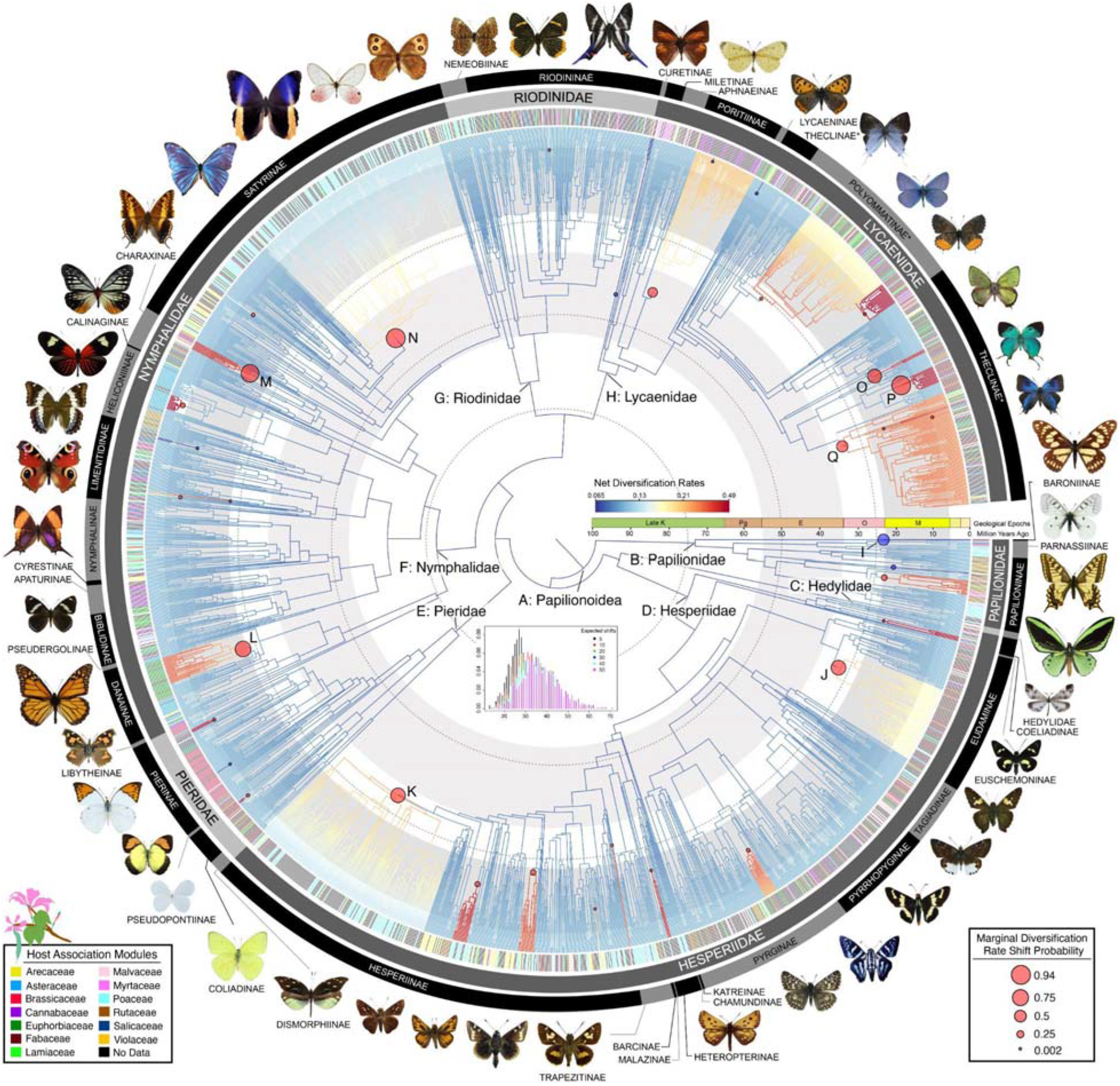
Evolutionary relationships and diversification patterns of butterflies. Time-calibrated tree based on 2,244 species, 391 loci, and 150 amino acid partitions. Branches show distinct shift configurations (circles) as estimated by clade-specific diversification models. Letters at nodes refer to clades with significant rate shifts (See section 6 of Supplementary Text). Colored lines in the outer ring beside tips indicate association with one of the 13 host modules (see section 17 of Extended Online Methods). Black lines in the host-association ring are species without data. The inset graph shows the posterior probability distribution, with alternative and expected number of diversification rate shifts, converging around 30.

To determine the geographic origin of butterflies, we used our dated phylogeny (Fig. 1) to conduct a global biogeographic analysis with 15,764 newly aggregated country-level distribution records (Table S4). Modeling with three different area categorizations, models of range evolution, and parameters (e.g., adjacency matrices, time slices, etc.) consistently recovered butterflies as originating in the Americas, in what is present-day western North America or Central America (Fig. 4, Table S5). All extant butterfly families excluding the Neotropical Hedylidae diversified ∼10–30 Ma after the Cretaceous Thermal Maximum (CTM), ∼90 Ma, when the global climate cooled by nearly 5° C^9^ (Figs. 1, 2). During the Cretaceous, butterflies appeared to be dispersing out of the Neotropics at a much higher rate than all other dispersal pathways (Fig. S2). As new butterfly lineages became established in other bioregions, other inter-bioregion dispersal avenues became more frequently used, particularly out of Indomalaya (Figs. S3, S4).

**Fig. 2.**
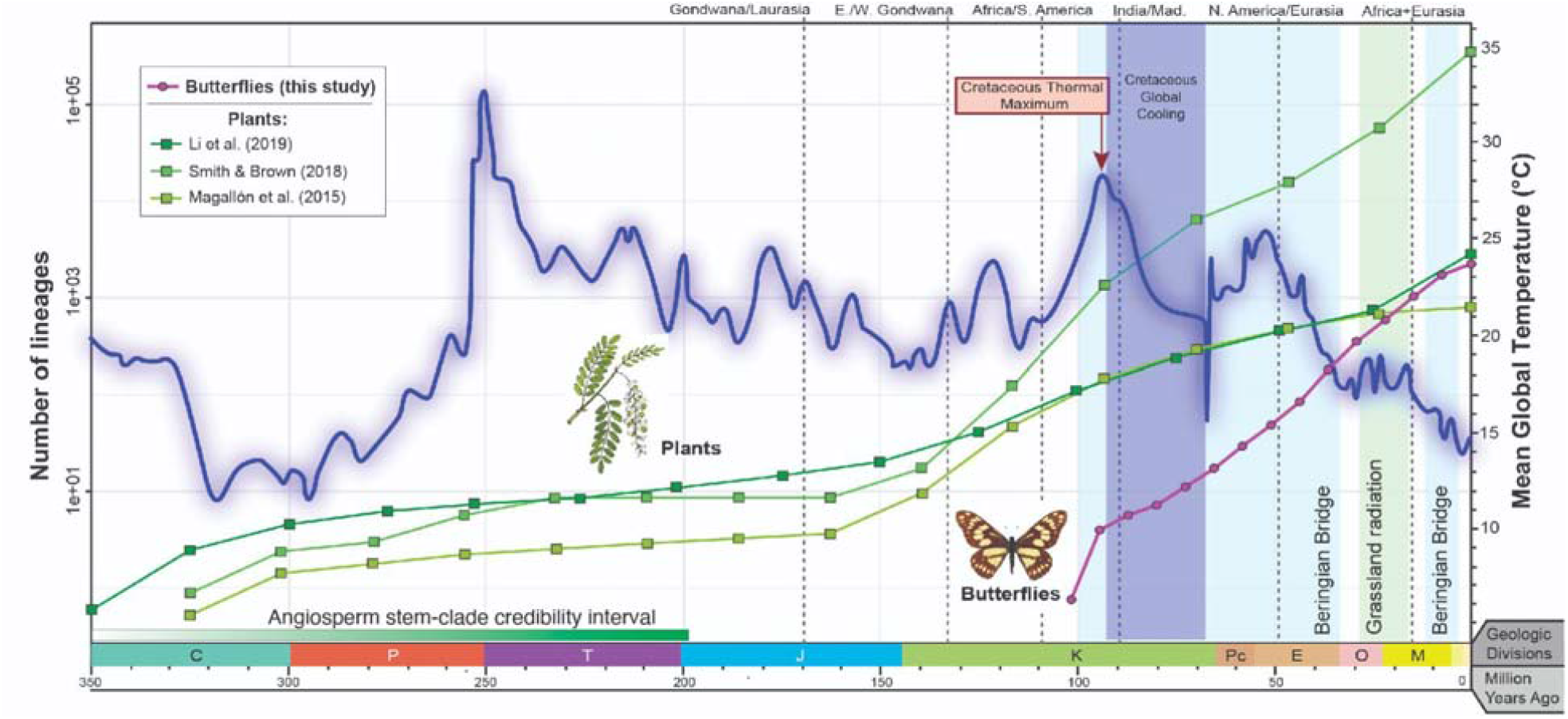
Global butterfly diversification over time. Butterfly diversity increased well after the origin of flowering plants. Colored lines with dots show butterfly diversity compared to vascular plant diversity from three widely accepted, recent plant diversification studies. Mean global temperature and major geological events during the last 350 Ma were calculated from Scotese et al.^85^. The angiosperm stem to clade credibility interval is a consensus of seven studies (see section 14 of Extended Online Methods).

Beginning around 60 Ma, the Neotropics served as an evolutionary “cradle”^10^ with high *in situ* butterfly speciation (Fig. S5), and many dispersal events occurring out of the Neotropics to other areas (Fig. S6). Relative rate of dispersal out of the Neotropics was still high during the early Cenozoic, although not as high as it was during the Cretaceous (Figs. S2, S3). Over the course of their evolution, butterflies experienced significantly higher speciation rates in the tropics compared to temperate zones (Data S1), and also more dispersal events out of the tropics (Fig. S6), as observed by high relative mean dispersal rates out of the tropics (e.g., tropical Indomalaya to temperate East Palearctic, and the Neotropics to Nearctic; Fig. 3). This pattern differs from that seen in mammals, which are thought to have dispersed primarily in the opposite direction during the Pliocene^11–13^. Some butterflies such as swallowtails showed greater immigration into the Neotropics, following a “museum” model of diversification (Fig. S7), corroborating prior findings^14^. The majority of dispersal events between the Neotropics and the Nearctic took place after the Eocene-Oligocene boundary, ∼33.9 Ma (Fig. S4). Two lineages dispersed from the East Palearctic around 17 Ma, and these appear to be the first colonizers of Europe: ancestors of a Nymphalini subclade including *Aglais, Nymphalis*, and *Polygonia*, and a clade of checkered skippers (Carcharodini; Table S6). Butterflies were present on what are now all modern continental landmasses by the late Eocene (Table S7).

**Fig. 3.**
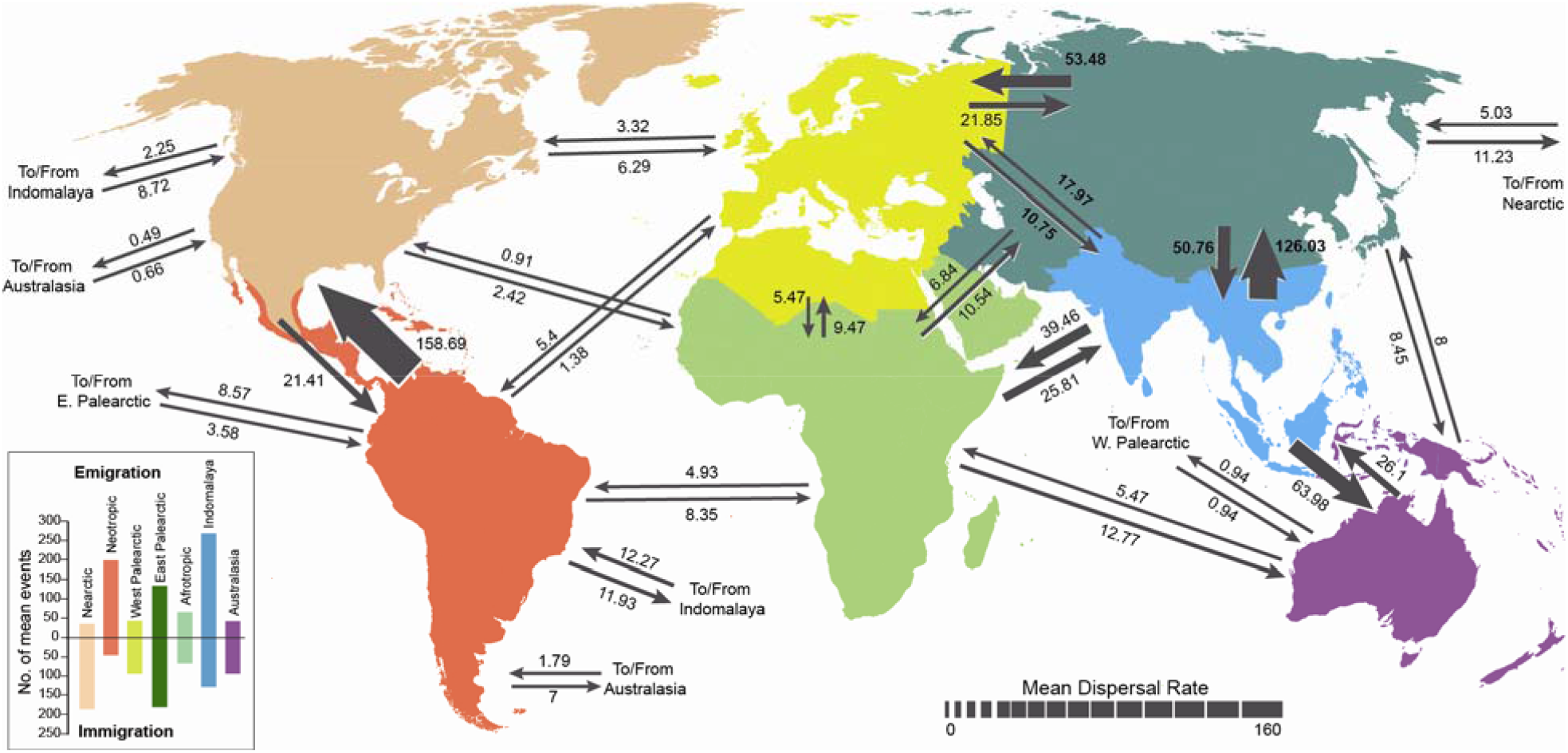
Relative mean dispersal rates of butterflies between different bioregions. Numbers associated with each arrow are the average rates from 1000 simulations using biogeographical stochastic mapping in BioGeoBEARS, which were then divided by 100 for ease of comparison (raw values can be found in Data S5).

**Fig. 4.**
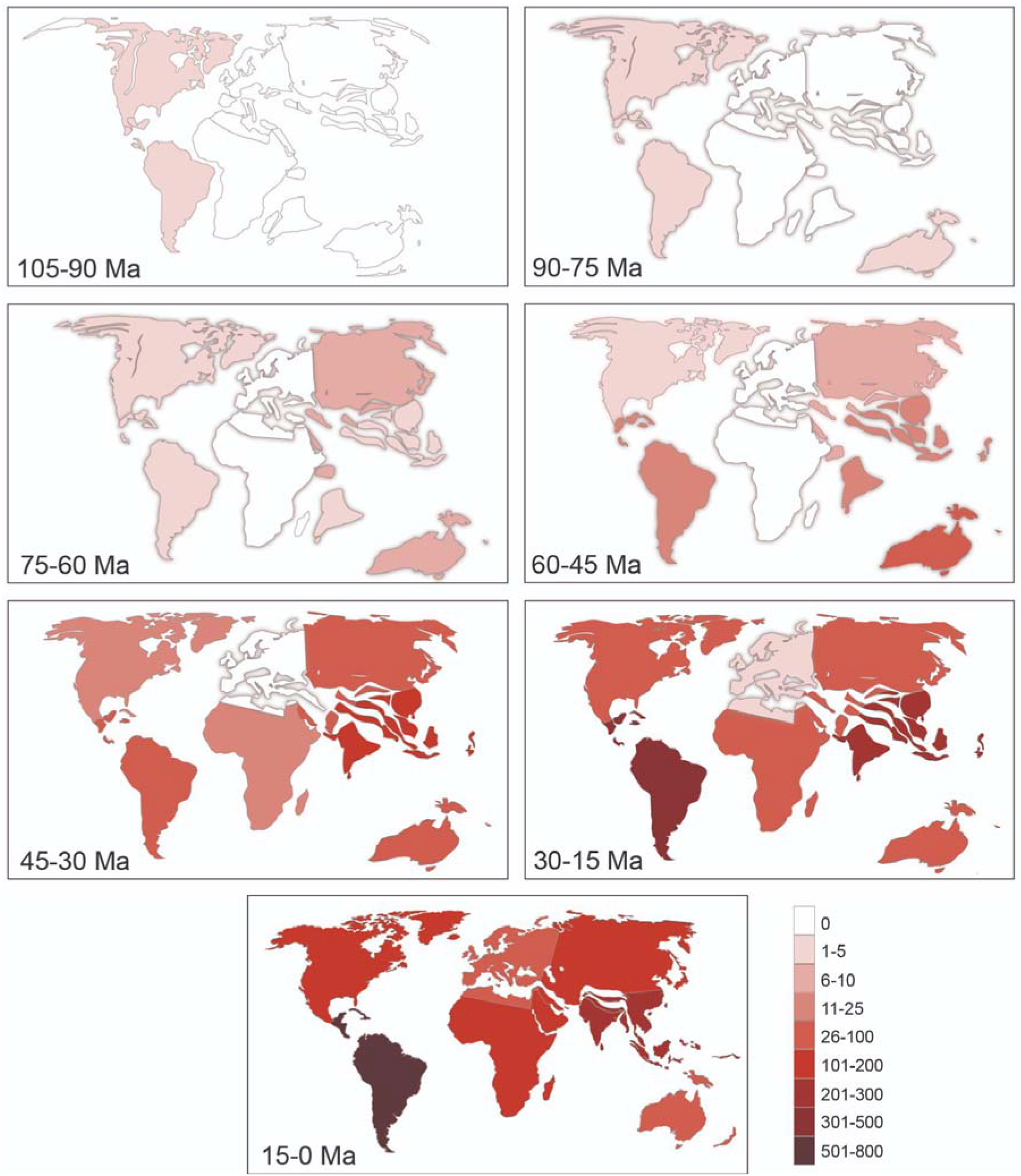
Distribution of ancestral butterflies over time. Each map corresponds to a 15 Ma interval of butterfly evolution. Bioregion color indicates the number of lineages in the Papilionoidea phylogeny that are associated with that bioregion during that time period, as determined by the BioGeoBears ASR.

The K/Pg boundary (∼66 Ma) marked a major global extinction event that dramatically reduced vertebrate and marine diversity^15^. Less is known, however, about the extent to which insects were affected by this event^16^. We used two analytical approaches to identify sudden extinction events and estimate changes in diversification rates, both of which concluded that butterflies, like amphibians and some mammals^17,18^, did not experience a major extinction event at the K/Pg boundary (Fig. S8). Plants experienced a vegetation turnover from gymnosperms to angiosperms in the Late Cretaceous^19^ and continued to increase in diversity up to the present (Fig. 2). The survival of many plant lineages across the K/Pg boundary may have buffered the effects of the mass extinction event of butterflies.

Two tests of extinction demonstrate that butterflies underwent two major extinctions after the K/Pg boundary — one at the Eocene–Oligocene transition (EOT) which coincides with the Eocene–Oligocene global extinction event (∼33.9 Ma), and another during the mid–Miocene that coincides with the mid-Miocene Climatic Optimum (∼14 Ma; Fig. S8). These were global cooling and warming periods, respectively, that led to the disappearance of many plant and animal lineages, followed by sharp floral and faunal turnover in temperate and tropical environments^15^. Speciation analyses with BioGeoBEARS^20^ (see Methods), also supports these findings; there was a dramatic drop in the number of speciation events around 34 Ma, especially in the East Palearctic region (Fig. S5), which occurred around the EOT extinction event that significantly reduced floral and faunal diversity of that region^21^. Soon thereafter, the families Hesperiidae and Nymphalidae experienced dramatic increases in the number of speciation events in the East Palearctic (Fig. S9). This marks a period when East Palaearctic habitats changed from dense forest to forest-steppe, temperate grassland, and mixed forest biomes^21^. These serve as primary habitats of the diverse modern butterfly groups – fritillaries, satyrs, and grassland skippers, which dominate this region today (Argynnini, Satyrini, and Hesperiinae, respectively).

It is often argued that butterfly evolution is closely tied to the diversification of angiosperms^5,6^. Yet, few studies have examined the timing and pattern of butterflies and their hosts on a broad, macroevolutionary scale. To understand the temporal diversification of butterflies and plants, we compiled 31,456 butterfly host records from 186 books and databases (Table S8). We examined how butterfly lineages increased through time and compared these results with four recent studies that examined plant diversification dynamics. We found that butterfly diversification lagged far behind the origin of angiosperms (Fig. 2), corroborating prior studies^7,22^. Ancestral state estimation provided support for Fabaceae as the host plant of the most recent common ancestor of butterflies (Tables S9-S10, Fig. S10), a widely accepted hypothesis^5^ that has lacked empirical support. The crown age of the most recent common ancestor of Fabaceae is thought to be ∼98 Ma^23,24^, largely coincident with the origin of butterflies.

Although the vast majority of butterfly larvae in our dataset are herbivores, a small number also feed on detritus, lichens, or insects (Table S8). The oldest associations in the entirely entomophagous Miletinae (Lycaenidae) appear to originate by 58.4 Ma (58.9–57.1 Ma), an estimate that corresponds with an earlier estimation of the origin of this group^25^ (Tables S3, S11). The Lycaenidae, with caterpillars that are ancestrally symbiotic with ants^7,26^, date back to 64.5 Ma (65.4–63.7 Ma) (Fig. S11), long after the origin of ants (139–158 Ma^27^). Together with plants, ants appear to have provided a template for diversification of Lycaenidae and certain members of its sister clade, Riodinidae.

To address the possibility that butterfly diversification may have been spurred by co-diversifying plants, we evaluated eight paleoenvironment-dependent models relating butterfly diversification to host-plant evolution^28^. We first estimated time-dependent variation of speciation and extinction rates with time-continuous birth-death models and compared them with constant-rate diversification models^29^. In the best-fitting model, speciation rates vary exponentially through time without extinction (92% of the trees, Akaike ω = 0.549). This time-dependent model is far better supported than the angiosperm-dependent models, the best of which was ranked fourth (Table S12). We also ran the HiSSE model^30^ as implemented in the R package hisse^31^ to test for a potential impact of host-plant preference on diversification dynamics. In total, we compared 18 different models of HiSSE and BiSSE-like implementations, accounting for hidden states to alleviate recent concerns regarding the reliability of SSE models and the high incidence of false positive results^32^. We found little or no indication that association with a specific plant group is linked to butterfly diversification, largely rejecting the hypothesis of hostplant driven diversification (Table S13, Data S2). These results indicate that coevolutionary interactions with angiosperm host-plants were unlikely to have been a driver of butterfly diversification at macroevolutionary scales. Even within families, hypotheses such as the correlated diversification of grasses with grass-feeding skippers (Hesperiinae) and satyrs (Satyrini)^33^ were uncorroborated (Table S13). Our results are consistent with smaller studies on particular butterfly groups such as skippers (e.g., Sahoo et al.^34^), that show that alternative drivers (hidden states) explain best butterfly diversification when compared to hostplant-dependent models.

Butterfly diversification may not have been driven by the availability of particular plants, but instead by host specificity (specializing on a few plants [e.g., Janz, Nylin and Nyblom^35^]). It has been proposed through the resource-use hypothesis^36^ that specialists that feed on few hosts have higher speciation and extinction rates. Numerous studies on vertebrates support this hypothesis^37,38^, but few studies have tested the hypothesis on insects. We examined host plant specificity on the butterfly phylogeny (Fig. 1) and found that more than two-thirds of butterfly species feed on a single plant family (67.7%), whereas less than one-third (32.3%) are generalists feeding on two or more (Table S14), a pattern largely in agreement with ecological studies on butterflies^39^. We also found that 94.2% of generalists feed on plant families that are significantly closely related compared to a randomly sampled null distribution, suggesting that generalists, although capable of feeding on different host families, still consume closely related plants. This finding supports the pattern reported by Ehrlich and Raven^6^ that related butterflies feed on related plants. Finally, we used trait-dependent methods to examine whether diversification rates changed following switches between generalist and specialist, and host plant shifts (Table S13). Our results demonstrate that – at least in analyses of Papilionoidea and Hesperiidae, where results were significant – specialists have lower rates of speciation (Fig. S12), contradicting the resource-use hypothesis.

This study includes the most comprehensive taxonomic sampling of butterflies to date to understand their evolutionary history. We also compiled two comprehensive trait datasets (geography and host plants) that demonstrate the importance of assembling large trait databases to test key hypotheses. Our study overturns some long-held assumptions of butterfly evolution and provides a framework for future studies of this model insect lineage. The consistency of results obtained using different approaches for each of our analyses suggests that our conclusions are robust. Assembling species-level datasets will be an important future goal to address fine-scale trends within particular butterfly clades.

Our data support the hypothesis that butterflies originated in the Americas in the Late Cretaceous 100 million years after the origin of angiosperms, and that they first fed on legumes. They dispersed from the Americas to the East Palearctic likely across Beringia ∼75 Ma before diversifying in the Paleotropics. They appear to have been minimally affected by the K/Pg mass extinction. While some evidence points to a Nearctic origin, the evidence for Nearctic versus Neotropical origin is not strong – we therefore only tentatively conclude that a Laurasian origin is possible. Diversification of some lineages has been impacted by host associations, but an escape-and-radiate model of coevolution with angiosperms does not appear to have been a powerful driver of diversification at a broad macroevolutionary scale. Butterfly evolution is better viewed as compatible with a model of diffuse coevolution^40^ in which plants provided an ecological template on which butterflies diversified as opportunities arose.

## Methods

### Taxon sampling and sequence acquisition

A total of 2,248 butterfly specimens representing 2,244 species in 1,644 genera was included for the molecular component of this study (Table S15). We obtained marker loci used for phylogenetic analysis via Anchored Hybrid Enrichment (AHE) exon capture of DNA extracts and subsequent Illumina sequencing^41^ or by bioinformatically extracting these sequences from published genomes and transcriptomes. We used the BUTTERFLY1.0 probe set^7^, which can capture up to 450 loci.

We extracted DNA from 1,915 specimens that were: 1) stored in ethanol and frozen; 2) dried and stored in glassine envelopes under ambient conditions (papered); or 3) dried, spread and pinned in a museum collection. Locus assembly and sequence clean-up followed the pipeline of Breinholt et al.^42^. Published sequences comprised: 1) genome assemblies; 2) genomic reads; and 3) paired or 4) single-end transcriptomes. Three sequence datasets were created for this study: nt123 (a nucleotide dataset with all codon positions), degen (a nucleotide dataset that excludes all synonymous changes, created using the perl script, Degen1 v.1.4^43,44^), and aa (an amino acid dataset translated from the nt123 dataset) (Data S3).

### Phylogenetic analysis and dating

Maximum likelihood (ML) tree inference was conducted on all three datasets in IQ-TREE 2.0 (nt123, degen, and aa), and parameter settings for each analysis can be found in Table S16. Branch support was calculated with 1,000 ultrafast bootstrap replicates (UFBS; ‘-B 1000’ command)^45,46^, and Shimodaira-Hasegawa approximate likelihood ratio tests (SH-aLRT; ‘-alrt 1000’ command)^47^. Quartet sampling (see *Section 8. Sequence quality check*) was performed on the most-likely degen359 and aa154 trees to examine the topology and provide more comprehensive and specific information on branch support. Four-cluster likelihood mapping (FcLM) analyses^48^ were performed on the degen and aa datasets to assess the placement of particular butterfly clades that have been the subject of previous phylogenetic studies. We applied this approach in addition to standard branch support metrics because the latter can be subject to inflated estimates^48^.

We obtained divergence time estimates using a penalized-likelihood based approach implemented in treePL^49^. We implemented three different methods for calibrating the trees to assess similarity among results. Method 1: Dating with secondary calibrations only. We used the 95% credibility intervals of Lepidoptera ages from Figure S12 of Kawahara et al.^50^ to assign minimum and maximum ages to 27 ingroup and 6 outgroup nodes in our tree. Method 2: Dating with fossils and one secondary root calibration. In this approach, we followed the guidelines of Parham et al.^51^ by calibrating nodes with 11 butterfly fossils that could be assigned to a butterfly lineage’s geological age with confidence as verified by de Jong^52^. None of the outgroup nodes could be calibrated because the only reliable fossils associated with our non-butterfly Lepidoptera were too young to influence the ages of the deeper nodes representing multi-superfamily clades. Consequently, preliminary treePL analyses yielded highly dubious age estimates for deep nodes on the tree, hundreds of millions of years older than expected based on the literature. We therefore added a single secondary calibration to the root of the tree. Although combining secondary and fossil calibrations in a single analysis can create redundancy that negatively impacts the resulting age estimates^53^, the limited fossil record of Lepidoptera made it a necessity in order to obtain comparable results derived primarily from fossils. We ran two versions of this method, each with a different root calibration. Method 2A used a maximum-age estimate of 139.4 Ma, based on the angiosperm age estimate of Smith and Brown^54^. Method 2B used a more conservative maximum-age estimate of 251 Ma, based on the older end of the credibility interval for the age of angiosperms in Foster et al.^55^. Both of these calibrations were used under the assumption that butterflies diverged from their moth ancestors after their most frequent host-plants, angiosperms, were already present^56,57^. Method 3: Secondary calibrations and six fossils. In this approach, we combined the 33 secondary calibrations from Method 1 with six fossil calibrations, including some of the fossils used in Method 2. Fossils previously used to time-calibrate trees of Kawahara et al.^50^ were excluded from this analysis to avoid circularity and redundancy with secondary calibrations. Whenever possible, redundant fossil calibrations from Method 2 were replaced with calibrations from unrelated fossils that could be associated with a different node in the same clade.

### Diversification rate analyses

We performed a Bayesian analysis of macroevolutionary mixtures using the program BAMM v.1.10.4^58^ to detect shifts in diversification rates between clades. The reversible-jump markov chain Monte Carlo was run for 50 million generations and sampled every 50,000 generations. Priors were estimated with the R package BAMMtools v.2.1.6^59^ using the command ‘setBAMMpriors’. The tree was trimmed in Mesquite v.3.6^60^ to remove all outgroups. Six analyses were performed using different priors for expected numbers of shifts (5, 10, 20, 30, 40, and 50 shifts).

Because BAMM has been criticized for incorrectly modeling rate shifts on extinct lineages (i.e., extinct or non-sampled lineages inherit the ancestral diversification process and cannot experience subsequent diversification-rate shifts^61,62^), we conducted a lineage-specific birth-death shift analysis^63^ in RevBayes^64^. We also utilized the R package castor v.1.6.9^65^ to infer deterministic lineage through time (dLTT), pulled speciation rate (PSR), and pulled diversification rate (PDR) plots for Papilionoidea and six of its families, excluding Hedylidae. Finally, we performed diversification analyses in TreePar v.3.3^66^ and TESS v.2.1.0^67^ to determine relative likelihoods of models with different amounts of shifts in diversification and turnover rates.

We conducted a series of diversification analyses to evaluate whether there is a correlation between butterflies and plants. For this, we used HiSSE (Hidden State Speciation and Extinction) and a BiSSE-like (Binary State Speciation and Extinction) implementation of HiSSE^30^ in the R package hisse^31^. We pruned outgroups from the tree (aa154 dated tree, Strategy A) and compared 20 HiSSE models and BiSSE-like implementations of HiSSE. The BiSSE equivalent of HiSSE tests whether there are different diversification rates associated with the two host-plant use states. Other models were built in the HiSSE framework to test alternative combinations of presence or absence of hidden state and host-plant use associations while also considering different transition rate matrices, net turnover rates τi (speciation plus extinction: λi + μi), and extinction fractions εi (extinction divided by speciation: μi/λi) (Table S17). We tested whether diversification rates were linked to feeding (A) as a larval specialist or generalist (Table S18), (B) on Poales (Table S19) in Papilionoidea, Hesperiidae, and Nymphalidae, (C) on Fabales (Table S20) in Papilionoidea and Nymphalidae, (D) on Brassicales (Table S21) in all butterflies and Pieridae, (E) on Fagales (Table S22), (F) on Poaceae module (Table S23), (G) on Fabaceae module (Table S24), and (H) on Fabaceae in Eudaminae (Tables S13, S24). We compared these different models of HiSSE and BiSSE-like implementations to account for hidden states to alleviate recent concerns regarding the reliability of SSE models and the high incidence of false positive results^32^.

### Biogeographic analyses

To assess the role of geography on diversification, we first aggregated data from multiple sources to create a global butterfly checklist for each country. Primary data sources included: 1) the *Lepidoptera and other life forms* database (http://ftp.funet.fi/index/Tree_of_life/insecta/lepidoptera); 2) WikiSpecies (https://species.wikimedia.org); and 3) the type locality of each species or subspecies in our list of valid butterfly names, which was obtained from 1, above. This initial global checklist was vetted using published country checklists and the ButterflyNet Trait Database (https://butterflytraits.org). Trait data from *ca*. 100 comprehensive and country-specific field guides have been entered into this database, allowing us to generate species lists to cross-validate checklists assembled^68^.

We estimated the ancestral area of origin and geographic range evolution for butterflies using two approaches: the ML approach of the DECX model^69^ as implemented in the C++ version^70,71^, available at https://github.com/champost/DECX), and with the program BioGeoBEARS v.1.1.2^20^. We designated 14 biogeographic regions across the globe (Fig. S13, Table S25), determined which of these regions were occupied by each species in our tree, and developed a 14-state character matrix. DECX uses a time-calibrated tree, the modern distribution of each species for a set of geographic areas, and a time-stratified geographic model that is represented by connectivity matrices for specified time intervals spanning the evolutionary history of clade of interest^72^.

Because we could not estimate immigration and emigration rates in DECX, we also ran BioGeoBEARS with 7 and 8 areas (Figs. S14, S15, Table S25). BioGeoBEARS analyses could not be run with 14 states due to computational limitations due to the complexity of our dataset (2,248 tree tips). The 7 and 8 bioregions largely correspond to the biogeographic realms defined by Udvardy^73^. In BioGeoBEARS, we implemented both Dispersal Extinction Cladogenesis (DEC) model^69,74^ and the Likelihood equivalent of the Dispersal-Vicariance approach (DIVALIKE)^75^ models and different adjacency matrices (Data S4). Both approaches gave largely consistent results, regardless of the model and parameters used (Tables S5, S26).

We performed biogeographic stochastic mapping to examine *in-situ* speciation, immigration, and emigration between the 7-bioregions in BioGeoBEARS. We followed the protocol of Li et al.^76^ with 1,000 simulations with the DEC model. Relative mean dispersal rates between all permutations of bioregions were calculated and presented in Figure 3 (see also Data S5). These mean dispersal rates represent dispersal of butterfly lineages throughout the entire evolutionary history of Papilionoidea, and thus cannot reveal changes in rates over time. In order to look at historical biogeography of butterflies during different epochs, rates along all possible inter-bioregion colonization rates were calculated at specific time intervals of 5 million years, following Li et al.^76^ (Table S27). These relative rates were then averaged to represent relevant geological time periods and presented in Figures S2-S4.

### Larval host plant analyses

Larval host records were compiled from numerous sources (Table S8, Data S6). Given the size of our host datasets and the scale of our analyses, we chose to examine relationships between individual butterfly species and host families that are consumed by their larvae. Plant families are commonly adopted as the taxonomic rank used for examining the evolution of host use^77,78^. For each plant-feeding butterfly species in our tree, we quantified host-plant richness and phylogenetic distance using six different metrics implemented in the package picante v. 1.8.2^79^. To calculate these metrics, we used the calibrated tree of seed plants from Smith and Brown^54^.

Because the number of host groups was too large for an ancestral state reconstruction (nearly 50 host-plant orders, ∼200 plant families plus insect associations), we first reduced the number of host groups by using a network analysis. The Beckett algorithm^80^, as implemented in the function ‘computeModules’ from the package bipartite^81^ in R v. 3.6.2^82^, assigns plants and butterflies to modules and computes the modularity index, Q. By maximizing Q, the algorithm finds groups of butterflies and hosts that interact more with each other than with other taxa in the network. Thus, host-plants that are assigned to the same module tend to be used by the same butterflies. We found 13 modules for butterfly host associations in our module analysis (Table S28, Table S17). We then conducted three larval host ancestral state reconstruction analyses using stochastic character mapping with SIMMAP in phytools v.0.7-70^83^ using the ‘make.simmap’ command. We reconstructed the ancestral state of (A) generalist versus specialist feeding (2 states, Data S7), (B) plant, lichen, Hemiptera, or Hymenoptera as a food source (4 states, Data S8), and (C) plant module (13 states as described above, Data S9).

We examined the speciation rate of butterflies over time with maximum-likelihood birth-death models to fit constant-rate (CST), time-dependent (TimeVar), and angiosperm-dependent (AngioVar) models of diversification using the R package RPANDA v.1.9^84^. These models jointly tested whether the diversification of angiosperms could have fostered the diversification of butterflies in a single statistical framework. We conducted a series of diversification analyses to evaluate whether there is a correlation between butterflies and plants. For this, we used HiSSE (Hidden State Speciation and Extinction) and a BiSSE-like (Binary State Speciation and Extinction) implementation of HiSSE in the R package hisse^30,31^.

## Acknowledgments

We thank M. Kuhn and E. Mavrodiev for assembling trait data. S. Epstein, T. Girard-Ang, P. Pezzi, and L. Xiao assisted with lab work. J. Barber, M. Brownlee, S. Cinel, J. Daniels, R. Godfrey, H. Gough, C. Hamilton, G. Hill, P. Houlihan, C. Huang, J. Miller, K. Miner, C. Mitter, K. Mitter, A. Renevier-Faure, J. Rubin, M. Scallion, Y. Sondhi, and L. Wu provided specimens or helped improve the manuscript. RAPiD Genomics (Gainesville, FL, USA) conducted sequencing. K. Meusemann (1KITE) provided unpublished FcLM scripts. T. Barve, C. Couch, H. Dansby, K. Casarella, R. Merritt, and X. Zheng helped create figures. High performance clusters at Brigham Young University, Smithsonian Institution, University of Florida, and Zoological Research Museum Alexander Koenig provided computational support.

## Data and materials availability

All data in the main text, supplementary materials, or in gzipped data archives will be made available at the Dryad Digital Repository (www.datadryad.org.xxxx) upon acceptance.

## Author contributions

Analysis: APSC, AYK, CE, CS, DMP, EAE, EFAT, FLC, HLO, JWB, ME, MPB, PBF, RAS, XL

Conceptualization: AYK, DJL, LR, JWB, ME, NEP, RG

Funding acquisition: AYK, DJL, MJZ, NEP, RG, WJ

Data assembly: CS, DJL, DMP, EAE, EAL, FLC, HW, JH, JWB, LR, ME, MI, MN, NEP, RG, VB, VS, WAVM

Methodology: CW, EH, FTGS, HP, JLS, JRK, JWB

Project administration: AYK, DJL, NEP

Sampling: ABBM, APSC, AYK, DJL, EFAT, FH, GT, HPR, JIM, JWB, ME, MY, NEP, RAS, RV, PV, ZFF

Sequence workflow: JWB, ME

Specimen identification and preparation: ADW, AM, APSC, AYK, DJL, HPR, JIM, JVL, KMD, ME, MY, NOM, RAS, RM, SCM, TD, YLN, ZFF

Supervision: AYK, DJL, JWB, ME, NEP, RG

Taxonomy and curation: DJL, GL, RG, VB

Visualization: AM, APSC, AYK, DJL, FLC, XL

Writing – original draft: APSC, AYK, DJL, DMP

Writing – review & editing: All authors

